# Site-Specific Fluorescent Labeling via SpyTag-SpyCatcher for Rapid Hybridoma Screening in Semi-Solid Medium

**DOI:** 10.64898/2026.08.21.746134

**Authors:** Amber Guo, Moris Wei, Jonny Wu, Xiaoxiao Li, Bruce Jiang

**Affiliations:** Shanghai Epizyme Biomedical, Shanghai, China

**Keywords:** hybridoma, monoclonal antibody, semi-solid medium, SpyTag-SpyCatcher, site-specific labeling, epitope preservation

## Abstract

Hybridoma screening in semi-solid medium typically employs antigens labeled with visible fluorophores (e.g., FITC, AF488) to enable single-step identification of antibody-secreting clones. However, conventional chemical conjugation via NHS-esters or isothiocyanate groups frequently modifies lysine residues located within epitopes, potentially abrogating antibody recognition of these critical regions. Here, we describe a SpyTag–SpyCatcher-based site-specific labeling strategy that circumvents epitope damage during semi-solid medium screening. A 16-amino-acid SpyTag was genetically fused to the C-terminus of the target antigen, enabling covalent conjugation to an sfGFP–SpyCatcher fluorescent probe. In semi-solid medium supplemented with SpyTag-antigen and sfGFP–SpyCatcher, positive hybridoma clones were readily identified by distinct fluorescent halos, whereas negative clones showed no detectable signal. Notably, the site-specific method yielded a significantly higher frequency of fluorescence-positive clones compared to the conventional AF488-labeled antigen method, suggesting that epitope preservation enhances screening recovery. Furthermore, this approach did not impair hybridoma growth or final clone positivity, offering a simple, rapid, and epitope-compatible method for monoclonal antibody screening.

## Introduction

The generation of monoclonal antibodies (mAbs) via hybridoma technology remains a cornerstone of biomedical research and therapeutic development. Semi-solid medium-based screening has emerged as a powerful approach for the rapid isolation of antigen-specific hybridoma clones, as it allows the simultaneous assessment of clone viability and antibody secretion within a single culture vessel^1,2^. In this system, fluorophore-labeled antigens—commonly conjugated via NHS-ester or FITC chemistry—are incorporated into the medium, enabling the visualization of antibody–antigen immune complexes as fluorescent halos surrounding positive clones^3^. This single-step visualization dramatically accelerates the screening process compared to traditional limiting dilution followed by multi-step ELISA^4^. Despite its convenience, chemical fluorophore conjugation poses a significant risk to epitope integrity. NHS-esters and isothiocyanates react indiscriminately with primary amines, predominantly lysine residues and the N-terminus^5^. When lysine residues reside within or adjacent to antibody-binding epitopes, such modifications can alter the antigen’s conformation or charge distribution, effectively masking the epitope and precluding the isolation of antibodies directed against these regions^6^. Consequently, researchers may inadvertently miss therapeutically or diagnostically relevant antibody specificities.

To address this limitation, instrument manufacturers have explored alternative non-chemical labeling strategies. For example, Molecular Devices has described an approach in which the antigen is genetically fused to a human Fc domain, and a fluorophore-conjugated anti-human Fc antibody is used as the fluorescent reporter in semi-solid medium screening^7^. While this strategy avoids direct chemical modification of the antigen, it introduces several constraints: the Fc domain is large (∼25 kDa) and may alter antigen conformation or create unwanted immunodominant epitopes; the anti-human Fc antibody is itself typically labeled via NHS-ester chemistry, reintroducing batch-to-batch variability; and the antigen–Fc/anti-Fc interaction is non-covalent and reversible, risking signal dissociation during extended culture.

The SpyTag–SpyCatcher system offers an attractive solution that overcomes these limitations. A genetically encoded 16-amino-acid peptide (SpyTag) and its cognate protein partner (SpyCatcher) undergo rapid, irreversible isopeptide bond formation under physiological conditions^8,9^. By fusing SpyTag to the C-terminus of the antigen and employing an sfGFP–SpyCatcher fusion as the fluorescent reporter, we reasoned that site-specific, covalent labeling could be achieved without compromising epitope integrity and without the drawbacks of Fc-based detection. Here, we demonstrate that this strategy enables efficient fluorescent conjugation, robust halo formation around antibody-secreting hybridomas, and a significantly higher recovery of positive clones compared to conventional chemical labeling—while eliminating the risk of epitope destruction and the limitations of Fc-fusion approaches.

## Results

### Design and in vitro validation of site-specific fluorescent conjugation

We first constructed the recombinant components for the site-specific labeling system. The NUCB1 (sequence from V276-E450) was expressed in Escherichia coli BL21(DE3) as a C-terminal SpyTag fusion (NUCB1–SpyTag, ∼26 kDa), while sfGFP–SpyCatcher (∼44 kDa) was produced in Expi293F cells. Both proteins were purified to homogeneity via Ni-NTA affinity chromatography (Fig. 1B).

**Figure 1.**
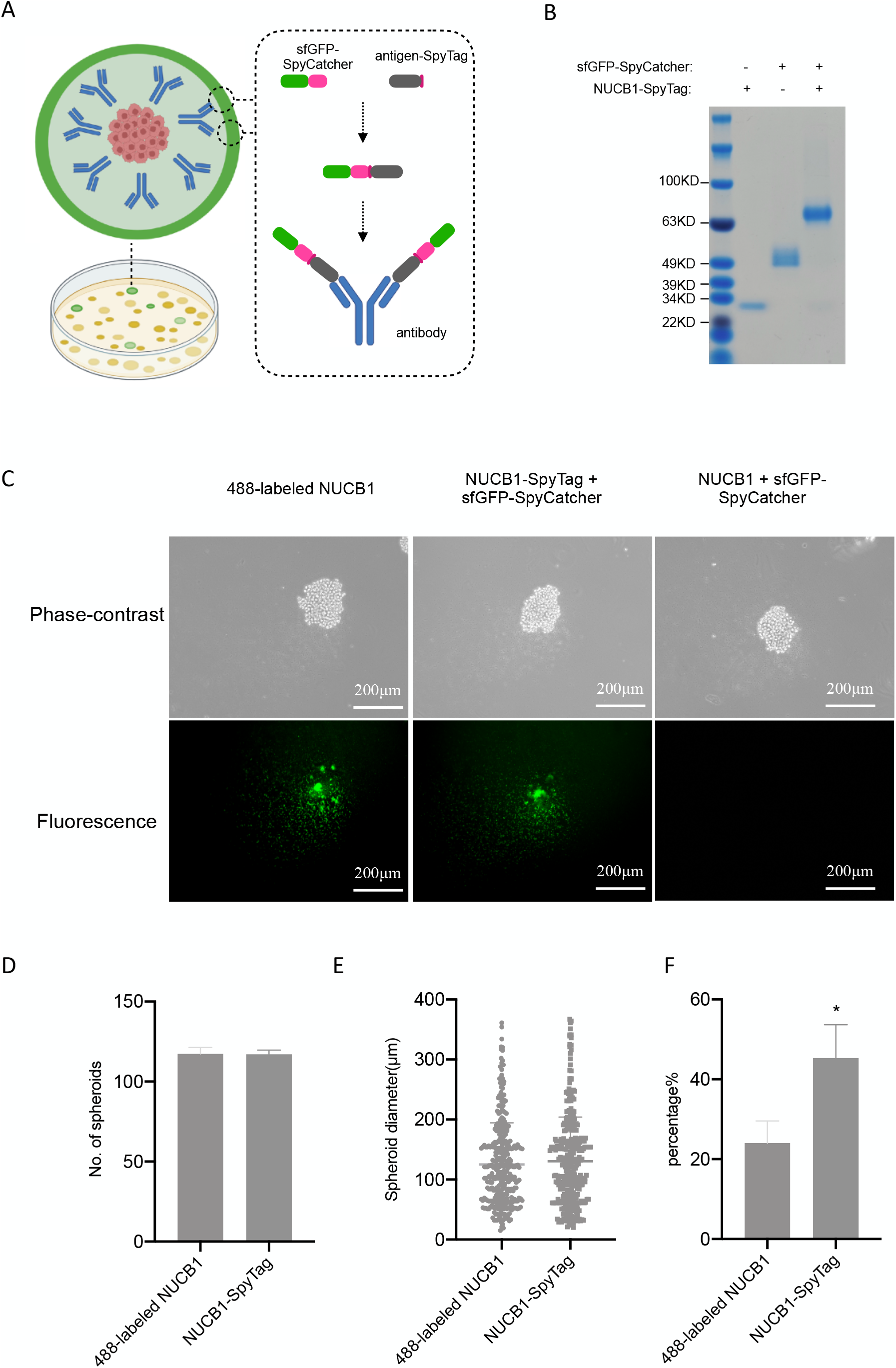
SpyTag–SpyCatcher site-specific fluorescent labeling for hybridoma screening in semi-solid medium. (A) Schematic illustration of the site-specific labeling strategy for screening. sfGFP– SpyCatcher (green) covalently conjugates to the C-terminal SpyTag of the antigen. Secreted antibodies from positive hybridoma clones bind the labeled antigen and form a localized fluorescent halo. Negative clones do not generate halos. (B) SDS-PAGE analysis of the conjugation reaction. (C) Representative phase-contrast (top row) and fluorescence (bottom row) images of hybridoma clones in semi-solid medium. (D) Clone formation efficiency in semi-solid medium following hybridoma fusion. The spheroids numbers in one well of 6-well plate was enumerated under the microscope. Data are mean ± SD; n = 3. (E) Statistics of spheroid diameter. ImageJ was used to determine the diameter. (F) Frequency of fluorescence-positive clones in the conventional and site-specific groups. Data are mean ± SD; n = 3; ^*^ indicates P < 0.05.

To assess conjugation efficiency, equimolar amounts of NUCB1–SpyTag and sfGFP–SpyCatcher were incubated at room temperature for 15 mins in 1×PBS. SDS-PAGE analysis revealed near-complete conversion to a higher-molecular-weight species at ∼70 kDa, consistent with the expected mass of the conjugated complex (Fig. 1B). These results confirm that the SpyTag– SpyCatcher reaction proceeds efficiently under conditions compatible with downstream screening applications, consistent with previous reports of rapid, high-yield conjugation in vitro^8,9^.

The operational principle of the screening platform is illustrated in Figure 1A. Upon mixing Antigen–SpyTag with sfGFP–SpyCatcher in semi-solid medium, spontaneous covalent conjugation generates the fluorescent antigen complex. When a hybridoma clone secretes antigen-specific antibodies, the secreted IgG binds the labeled antigen and forms an immune complex that remains localized around the clone, producing a visible fluorescent halo. Non-secreting clones or clones secreting irrelevant antibodies do not generate such halos.

### Performance of the SpyTag–SpyCatcher system in hybridoma screening

We next evaluated the site-specific labeling strategy in a functional hybridoma screening assay using an acquired hybridma cell-line secreting antibody against NUCB1. Three experimental groups were established: (i) the conventional method using chemically conjugated AF488-labeled antigen; (ii) the SpyTag–SpyCatcher site-specific method (NUCB1–SpyTag + sfGFP–SpyCatcher); and (iii) a negative control for the site-specific method using unmodified antigen lacking the SpyTag sequence (antigen + sfGFP–SpyCatcher).

As shown in Figure 1C, both the conventional AF488-labeled antigen and the SpyTag–SpyCatcher conjugate produced bright, well-defined fluorescent halos surrounding antibody-secreting hybridoma clones. The fluorescence signals were clearly detectable by standard fluorescence microscopy and were colocalized with cell clusters visible under phase-contrast imaging. In contrast, the negative control group (antigen without SpyTag) showed no specific fluorescence accumulation around clones, confirming that halo formation depends on the covalent SpyTag– SpyCatcher linkage and specific antibody–antigen binding.

To assess whether the site-specific labeling components affected hybridoma viability or proliferation, we compared clone formation efficiency between the conventional and SpyTag– SpyCatcher groups immediately following hybridoma fusion. Note that we applied a combinational strategy for antigen preparation, where Trx-NUCB1 for immunization and NUCB1 with/without the SpyTag at the C-termius for screening, in order to avoiding the contamination of antibody against the tag (Trx) during screening. No significant difference in the number of viable clones or the spheroids diameter was observed between the two groups (Fig. 1D,1E), indicating that the presence of NUCB1–SpyTag and sfGFP–SpyCatcher in the semi-solid medium does not impair hybridoma growth or colony formation.

We then quantified the frequency of fluorescence-positive clones in both screening conditions. Strikingly, the site-specific SpyTag–SpyCatcher group exhibited a significantly higher percentage of clones with distinct fluorescent halos compared to the conventional AF488-labeled antigen group (Fig. 1F). This finding suggests that chemical modification of the antigen by NHS-ester conjugation partially masks or destroys antibody-binding epitopes, thereby reducing the recovery of antigen-specific hybridomas. In contrast, the site-specific labeling strategy preserves epitope integrity, enabling the detection of a broader repertoire of antibody-secreting clones. Finally, to validate that fluorescence-positive clones identified by the SpyTag–SpyCatcher method were true antibody secretors, we randomly picked 156 halo-positive clones into 96-well plates and subjected them to confirmatory ELISA. All survived clones were confirmed positive for antigen-specific antibody secretion (data not shown), indicating that the fluorescent halos observed in the semi-solid medium are a faithful predictor of specific antibody production.

## Discussion

We have developed and validated a SpyTag–SpyCatcher-based site-specific fluorescent labeling strategy for hybridoma screening in semi-solid medium. This approach addresses a critical limitation of conventional chemical conjugation methods: the indiscriminate modification of lysine residues that can destroy or mask antibody-binding epitopes. By genetically fusing a 16-amino-acid SpyTag to the C-terminus of the antigen and employing an sfGFP–SpyCatcher fluorescent probe, we achieved rapid, covalent, and site-specific labeling. The resulting fluorescent antigen complex enabled the direct visualization of antibody-secreting hybridoma clones and, notably, yielded a significantly higher frequency of positive clones compared to conventional AF488-labeled antigen.

The preservation of epitope integrity represents the principal advantage of this method. In conventional NHS-ester or FITC labeling, the degree and distribution of fluorophore attachment are stochastic and difficult to control^53^. Epitopes rich in lysine residues—common in many immunodominant regions—are particularly vulnerable to such modifications. The SpyTag– SpyCatcher system confines the conjugation event to a single, defined site at the C-terminus, leaving the remainder of the antigen surface unaltered. This feature is especially valuable when screening for antibodies against conformational epitopes or when the antigen’s immunogenic profile is unknown.

The most compelling evidence for epitope preservation in our study is the quantitative difference in screening recovery: the site-specific method consistently identified more fluorescence-positive hybridoma clones than the chemically labeled antigen. We interpret this difference as a direct consequence of epitope masking or destruction by random AF488 conjugation. When lysine residues within or near antibody-binding sites are modified, the corresponding epitopes become invisible to the hybridoma repertoire, causing specific clones to be scored as negative. Because the SpyTag–SpyCatcher method leaves these surfaces untouched, it effectively expands the detectable epitope landscape and increases the yield of positive clones from a single fusion. Future study may be needed to directly confirm this advantage of site-specific conjugation e.g. designing an antigen peptide enriching lysine.

Recently, instrument manufacturers have introduced alternative non-chemical labeling strategies for semi-solid medium screening. Molecular Devices has described an approach in which the antigen is genetically fused to a human Fc domain, and a fluorophore-conjugated anti-human Fc antibody is used as the fluorescent reporter^7^. While this strategy avoids direct chemical modification of the antigen, it introduces several limitations that are overcome by the SpyTag– SpyCatcher system. First, the Fc domain is large (∼25 kDa) and may alter the native conformation of the antigen, affect solubility, or create unwanted immunodominant epitopes that compete with the target antigen for antibody binding^10^. In contrast, SpyTag comprises only 16 amino acids and is unlikely to perturb the structure or function of the fused protein8. Second, the anti-human Fc antibody is itself typically labeled via NHS-ester chemistry, reintroducing the risk of batch-to-batch variability, compromised binding affinity, and unconjugated dye contamination^7^. The sfGFP– SpyCatcher fusion used here is a recombinant protein with genetically encoded 1:1 fluorophore stoichiometry, eliminating chemical conjugation entirely. Third, the antigen–Fc/anti-Fc interaction is non-covalent and reversible; during the extended incubation in semi-solid medium (typically 10– 14 days), dissociation of the fluorescent reporter may reduce signal intensity and compromise discrimination of low-secreting clones. The SpyTag–SpyCatcher interaction forms a spontaneous, irreversible covalent isopeptide bond that remains stable throughout the culture period^8,9^. Fourth, human Fc may bind to Fc receptors on hybridoma or feeder cells, or to Fc-binding proteins present in serum-supplemented medium, generating background fluorescence and false-positive signals^11^. SpyTag has no known biological activity or off-target binding in mammalian cell culture. Our results demonstrate that the SpyTag–SpyCatcher components are fully compatible with the semi-solid medium environment. The conjugation reaction proceeded efficiently under culture conditions, and neither the SpyTag-fused antigen nor the sfGFP–SpyCatcher fusion protein exhibited detectable toxicity toward hybridoma cells. The equivalent clone formation efficiency between the site-specific and conventional methods confirms that the increased recovery of positive clones is attributable to improved antigen quality rather than enhanced cell growth. The modular architecture of this platform offers additional practical benefits. The SpyTag peptide is small and can be readily appended to either terminus of virtually any recombinant antigen via standard molecular cloning^9^. The SpyCatcher module can likewise be fused to alternative fluorophores (e.g., mCherry, iRFP) or other functional reporters, enabling multiplexed screening or adaptation to different detection systems^2^. Furthermore, because both components are recombinantly expressed, the method ensures batch-to-batch consistency that is often challenging to achieve with chemically conjugated reagents.

One consideration is that this approach requires recombinant expression of both the antigen– SpyTag fusion and the fluorescent SpyCatcher probe, which may represent a modest upfront investment compared to commercial chemical labeling kits. However, given the high conjugation efficiency, the increased yield of positive clones, and the ability to generate large, homogeneous batches of labeled antigen, this investment is readily justified for projects where epitope preservation is paramount. In cases where the C-terminus of the antigen is buried within the protein core or essential for function, an N-terminal SpyTag fusion could be explored as an alternative configuration^8^.

In summary, the SpyTag–SpyCatcher site-specific labeling strategy provides a robust, epitope-compatible alternative to both chemical fluorophore conjugation and Fc-fusion-based detection for hybridoma screening in semi-solid medium. By preserving epitope integrity and thereby increasing the recovery of antigen-specific clones, this method enhances the probability of isolating antibodies against the full repertoire of antigenic determinants.

## Materials and Methods

### Protein expression and purification

The gene encoding NUCB1–SpyTag or NUCB1 was cloned into pET28a and expressed in Escherichia coli BL21(DE3) with a N-termial 6×His tag. Cells were grown in LB at 37°C to an OD_600_ of 0.6, induced with 1mM IPTG overnight at 16°C, and harvested by centrifugation. The protein was purified via Ni-NTA affinity chromatography followed by size-exclusion chromatography in PBS.

The gene encoding sfGFP–SpyCatcher was cloned into pcDNA3.1 and transiently transfected into Expi293F cells using PEI. The culture medium Transpro CD1 (Duoning, MS002-500ml-02) was used. After 1 week culture at 37°C with 80 rmp/min, the supernatant was harvested and the protein was purified via Ni-NTA affinity chromatography followed by size-exclusion chromatography in PBS.

### SpyTag–SpyCatcher conjugation and SDS-PAGE analysis

Purified NUCB1–SpyTag and sfGFP–SpyCatcher were mixed at a 1:1 in PBS and incubated at room temperature (22-25°C) for 15min. The reaction mixture was analyzed by SDS-PAGE on a 12% gel under reducing conditions and visualized by commassie-blue staining.

### Chemical labeling of antigen (conventional method)

NUCB1 was labeled with AF488 NHS-ester according to the manufacturer’s instructions (Epizymem, JQ101). Briefly, 100 uM NUCB1 was reacted with a 10 fold molar excess of AF488 NHS-ester in PBS for 1h at 37°C. Unreacted dye was removed by the desalting column.

### Hybridoma generation and semi-solid medium screening

BALB/c mice was immunized with the antigen as the previously described^13^. Spleenocytes from immunized mouse were fused with SP2/0 myeloma cells using PEG1450. Fused cells were immediately resuspended in semi-solid medium (Epyzime, RZ201). For the conventional group, AF488-labeled NUCB1 was added at 80 nM. For the site-specific group, NUCB1–SpyTag or NUCB1 and sfGFP–SpyCatcher were added at 80 nM (molecular ratio: 1 :1). A total of 1 ×10^6^ cells (calculated by SP2/0) were seeded at one well in 6-well plates and incubated at 37°C, 5%CO_2_ for 8 days.

### Imaging and clone enumeration

Fluorescence and phase-contrast images were acquired using Olympus CKS53 microscope equipped with 10× objective, GFP filter sets. Clone formation was assessed by counting visible colonies per well under phase contrast. Fluorescence-positive clones were identified by the presence of a distinct fluorescent halo surrounding the cell cluster.

### ELISA confirmation

Halo-positive clones were picked and expanded in DMEM supplemented with 10% FBS. Culture supernatants were tested for antigen-specific antibody by indirect ELISA. NUC1 was coated onto elisa plate at 5ug/mL. After blocking with 3% BSA, supernatants were added and incubated for 1h at 37°C. Bound antibodies were detected with HRP-conjugated anti-mouse secondary antibody (Epizyme, LF101) and developed with TMB (Epizyme, HJ000). Absorbance was measured at 450nm using a plate reader.

### Statistical analysis

Data are presented as mean ± standard deviation (SD) from three independent experiments. Statistical significance was determined by unpaired two-tailed Student’s t-test using Graphpad prism7. A P value < 0.05 was considered statistically significant.

## Contribution

Conceptulization: Bruce Jiang, Amber Guo;

Experiment design: Bruce Jiang, Amber Guo;

Perform the experiment: Amber Guo, Moris Wei, Xiaoxiao Li;

Data curation: Bruce Jiang, Amber Guo.

Draft writing: Bruce Jiang;

All authors discussed the data and help the revision of the manuscript.

